# Feedback training can increase detection of happiness in ambiguous facial expressions in children and adults with autism spectrum disorders

**DOI:** 10.1101/524322

## Abstract

Recognition of subtle emotional facial expressions is challenging for some individuals with autism spectrum disorders (ASD). Training that targets recognition of low intensity emotional expressions may therefore be effective as an intervention to improve social-emotional skills. This paper reports the results of two randomised controlled experiments looking at the effect of a training methodology designed to increase the recognition of happy emotion in low intensity happy facial expressions. The first study implements this training with children with autism spectrum disorder (ASD, N = 14) and the second study implements this training with adults with ASD (N = 27). The training paradigm used images from a morph sequence that mixed a happy expression with a mixed-emotion ‘norm’ expression to create a sequence of varying intensity happy expressions. Participants were asked to say whether or not individual faces from the sequence were happy, to measure their happiness detection threshold. Participants that received active training were given biased feedback to shift their detection threshold, while participants that received control training were given feedback consistent with their baseline threshold. There was some statistical evidence that thresholds in the active training group shifted more than in the control group. This suggests training was successful in increasing the number of expressions that individuals identified as happy. However, there was no evidence that training increased facial expression recognition accuracy, as measured by the Reading the Mind in the Eyes task completed after training (Study 2).

## Introduction

There is evidence for reduced perceptual sensitivity for certain facial expressions (Doi et al., 2013; Rump, Giovannelli, Minshew, & Strauss, 2009; Wallace et al., 2011) and possible biases forwards recognition of negative emotions in ASD (Evers, Steyaert, Noens, & Wagemans, 2015). Reduced accuracy in recognising low intensity emotional expression has been associated with poorer adaptive functioning in ASD (Wallace et al., 2011), while a bias towards detecting negative emotions in facial expressions has been associated with poorer communication and emotional intelligence in ASD (Eack, Mazefsky, & Minshew, 2015). Training that targets the interpretation of low intensity emotional expressions in order to increase accuracy and reduce bias may therefore be effective as an intervention to improve social-emotional skills in individuals with ASD.

Cognitive bias, including attentional bias and interpretation bias, have been proposed to be a causal mechanism in a number of psychiatric disorders which a lead to negative affect. Biases in the interpretation of facial expressions have been found in anxiety, depression and conduct disorder (Bell et al., 2011; Bourke, Douglas, & Porter, 2010; Schönenberg & Jusyte, 2014; Yoon, Yang, Chong, & Oh, 2014). Cognitive bias modification (CBM) procedures that target attentional or interpretation biases have been developed as a therapy for individuals with psychiatric disorders (Heeren, Mogoaşe, Philippot, & McNally, 2015). For example Amir et al. (2009) trained participants with social phobia to attend to neutral instead of disgusted faces using a dot-probe paradigm leading to reductions in anxiety symptoms. Similarly, Beard and Amir (2008) gave social anxious individuals feedback reinforcing the benign interpretations over threat related interpretations of ambiguous scenarios leading to a decrease in social anxiety symptoms.

Recently a CBM procedure has been developed which aims specifically to modify biases in the perception of ambiguous facial expressions (Dalili, Schofield-Toloza, Munafò, & Penton-Voak, 2016; Penton-Voak, Bate, Lewis, & Munafò, 2012; Penton-Voak et al., 2013). In this training, participants are presented, in a random order, with facial expressions from a 15-step morph sequence between a happy and angry (or sad) expression. They are asked to categorise each expression as being one of the two emotions. The point in the morph sequence at which each participant switches from one emotion category to another is determined at baseline. In training participants then categorise the expressions again, this time receiving feedback about their accuracy after each response. Participants in the active training group receive feedback telling them that two more expressions just on the angry (or sad) side of their own category boundary (i.e., relatively ambiguous expressions that they characterized, on average as angry (or sad) at baseline) are actually happy. Participants in the control group receive feedback that is consistent with their baseline category boundary. Penton-Voak et al. (2013) trained youths at risk of criminal offending to categorise more expressions in a happy-angry morph sequence as happy. Immediately after training and two weeks later, those who received training to modify their category boundary showed reduced self-reported and observer-reported aggressive behaviour, compared to those who received control training. This suggested that this training, which targets the interpretation of low intensity expressions, can affect interpretation of emotions outside the lab and have a positive effect on mood and behaviour.

A possible reason for difficulty in recognising low intensity expressions in ASD is the presence of perceptual bias towards negative emotions. Recent studies which have examined confusion patterns in expression recognition tasks suggest that individuals with ASD show a bias towards certain negative expressions (Eack et al., 2015; Evers et al., 2015). In these studies participants with ASD were more likely to label neutral and happy expressions as negative emotions. Psychiatric disorders are common in ASD (Joshi et al., 2010), so it is possible that co-morbid psychiatric disorders, rather than ASD itself, may underlie these biases. Nonetheless, negative biases may contribute to problems with emotion recognition accuracy in ASD and, therefore, reducing negative biases may improve emotion recognition accuracy and social skills.

Here we investigated whether it was possible to increase detection of happiness in relatively ambiguous expressions of low intensity happiness. The training procedure was based on that used in previous studies (Dalili et al., 2016; Penton-Voak et al., 2012; Penton-Voak et al., 2013). However, instead of using stimuli from morph sequences that spanned two emotional expressions, the stimuli ranged from one expression, through an ambiguous expression (created by averaging 7 expressions), to the corresponding anti-expression (an expression in which the features have moved in the opposite direction from the emotional expression; for example, eyebrows are raised rather than lowered, see Method section for technical details). We chose to morph the emotional expressions with an ambiguous expression, rather than another emotional expression, because we wanted to increase the chances of improving recognition to one particular emotion without reducing recognition of another, given evidence of generally poor recognition of all emotional expressions in ASD (Harms, Martin, & Wallace, 2010). As the anti-expressions at the end of the sequence do not correspond strongly to a particular emotional state, reducing sensitivity to these anti-expressions should not impair emotion recognition ability. Although similar training procedures have been successfully employed previously (Penton-Voak et al., 2012; Penton-Voak et al., 2013), broad difficulties in face processing in ASD (Dawson, Webb, & McPartland, 2005) may make training with expression morphs unfeasible in this particular population. In the first experiment, we therefore sought to determine whether modification feedback (as opposed to control feedback) in training could increase the number of morph sequence expressions labelled as happy in a group of children and adolescents with ASD.

## Study 1

### Method

#### Participants

Eighteen children and adolescents (11-17 years) with ASD were recruited from an autism-specific educational setting that caters for individuals with mild to moderate intellectual difficulties. The school provided confirmation of diagnosis with reference to official statements of Educational Needs. All participants had moderate language abilities and normal or corrected-to-normal vision. Four participants did not complete the task, leaving a total of 14 participants, 7 who had been randomly assigned to the modification condition (1 female, mean age = 14 years, SD = 2 years), and 7 who had been randomly assigned to the control condition (1 female, mean age = 15 years, SD = 1 year). This study was approved by the University of Bristol Faculty of Science Research Ethics Committee.

#### Stimuli

Stimuli were 15 images from a happy to anti-happy morph sequence (see Figure 1). The sequence was created by morphing between a happy expression and an anti-happy expression to create a sequence in which the expression ranged from unambiguous intense happiness though low intensity happiness to a 30% anti-happy expression.

**Figure 1.**
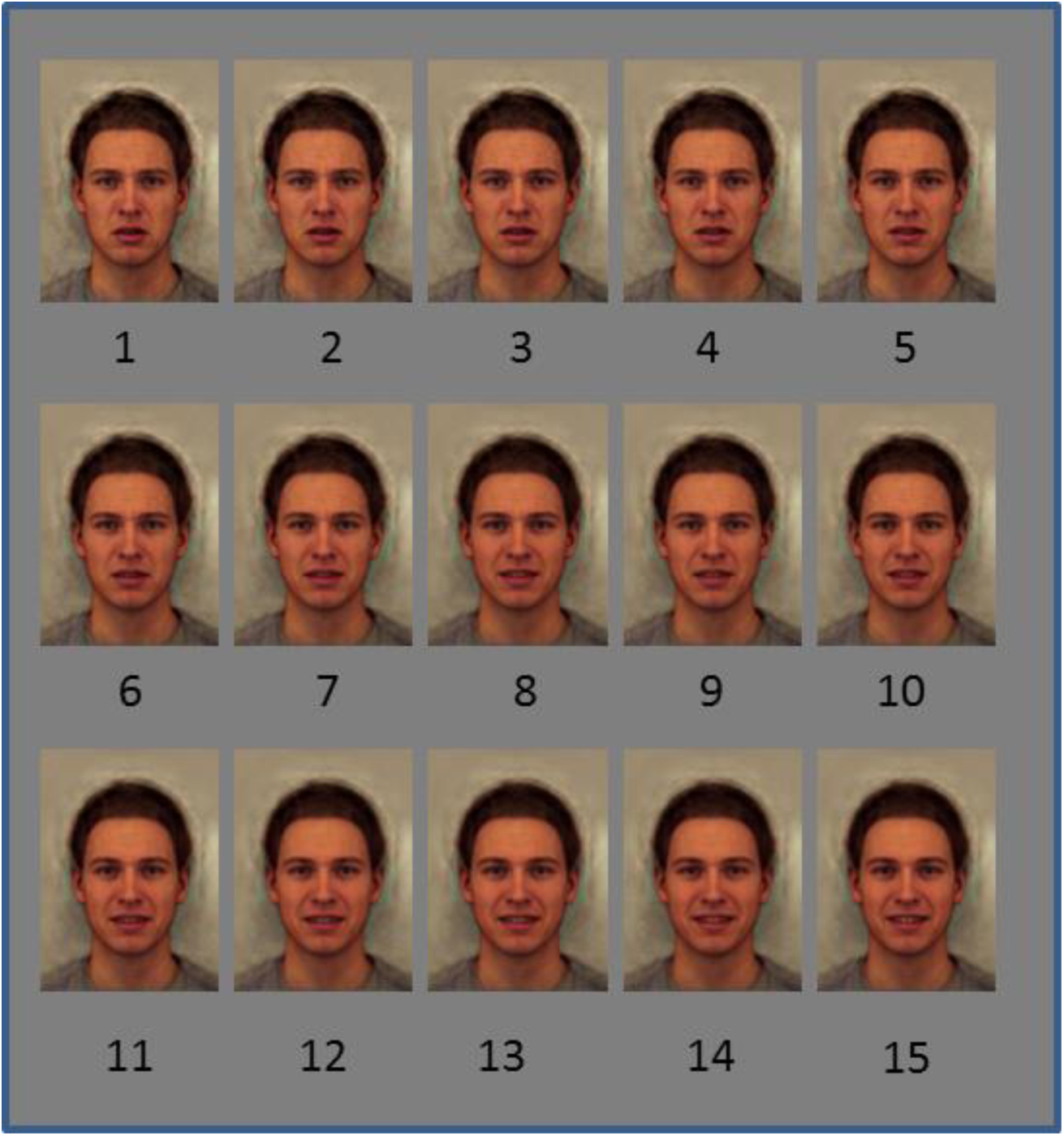
Morph sequence images 1-15

The happy expression was created using computer graphics techniques (Tiddeman, Burt, & Perrett, 2001) to average shape, colour and texture information from photos of 20 adult males with a happy expression taken from the Karolinska Directed Emotional Faces set (Lundqvist, Flykt, & Öhman, 1998). Additionally an emotionally ambiguous “norm” expression was created by averaging photos of the same 20 males with happy, sad, angry, surprised, scared, disgusted and neutral expressions.

A morph sequence was then created between the happy expression and the norm expression. This was then extended past the norm expression to create a 30% anti-happy expression. 15 equally spaced steps between the 100% happy expression and the 30% anti-expression were then selected. Therefore, moving along the sequence in Figure 1, the first 3 pictures contain components of the anti-expressions, while the next 12 pictures contain components of the veridical expression at increasing intensity levels. Including the 30% anti-expression at the end of the sequence, rather than stopping at the norm expression ensured that roughly half the morph sequence pictures were categorised as the target emotion at baseline. This was to keep the training consistent with that employed in previous studies (Penton-Voak et al., 2012; Penton-Voak et al., 2013) to stop a bias developing simply because of the predominance of one response at the outset.

#### Procedure

Participants in the modification condition and the control condition all completed three experimental phases: baseline, feedback and test. In each phase, participants saw the 15 images presented in a random order and judged whether each was happy or not. In the baseline and test phases each image was shown 3 times, giving a total of 45 trials. In the feedback phase there were 6 blocks of 31 trials in which images 1-2 and 14-15 were presented once, images 3-5 and 11-13 were presented twice and images 6-10 were presented three times. This was to reduce the length of training and to focus training on the critical frames in the morph sequence (where the threshold was expected to lie on the basis of pilot data). The feedback phase lasted around 20 minutes with the baseline and test phases occurring immediately before and after the feedback phase respectively.

The proportion of faces each participant judged to be happy at baseline was used to select baseline recognition threshold, corresponding to a particular frame from the morph sequence. In the feedback phase, participants in the control condition received feedback consistent with their baseline threshold, whereas participants in the modification condition received feedback that attempted to shift their baseline threshold by two frames, so an extra two frames just below threshold were categorised as happy. This two-step change is the same as in previous training (Penton-Voak et al., 2012; Penton-Voak et al., 2013).

In each trial, participants were presented with a central fixation cross for 1500–2500 ms (randomly jittered), followed by a face (562 by 762 pixels) for 150 ms. A mask of visual noise was then presented for 150 ms, after which a central question mark was displayed until participants responded by pressing one of two keys on the keyboard. In feedback blocks, responses were followed by a screen, displayed for 1000 ms, with the text ‘Correct (Incorrect)! That face was happy (not happy)’. A blank screen appeared between trials until participants pressed any key to continue. Most participants were able to press the response keys themselves. However, if a child had difficulty initiating the motor response, they responded verbally and the experimenter pressed the corresponding response key.

#### Analysis

Change in threshold from baseline to test phase was used to assess the effectiveness of modification feedback compared to control feedback in altering categorisation. Lower thresholds reflect a greater number of faces categorised as ‘happy’. Thresholds were analysed with a mixed model ANOVA with time (baseline, test) as a within-participant factor, and condition (modification, control) as a between-participant factor.

## Results

The mean thresholds at baseline were 9.14 (*SD* = 2.67) frames in the modification condition and 10.29 (*SD* = 1.11) frames in the control condition. The mean thresholds at test were 7.86 (*SD* = 1.95) frames in the modification condition and 10.00 (*SD* = 1.53) frames in the control condition (see Figure 2).

**Figure 0.**
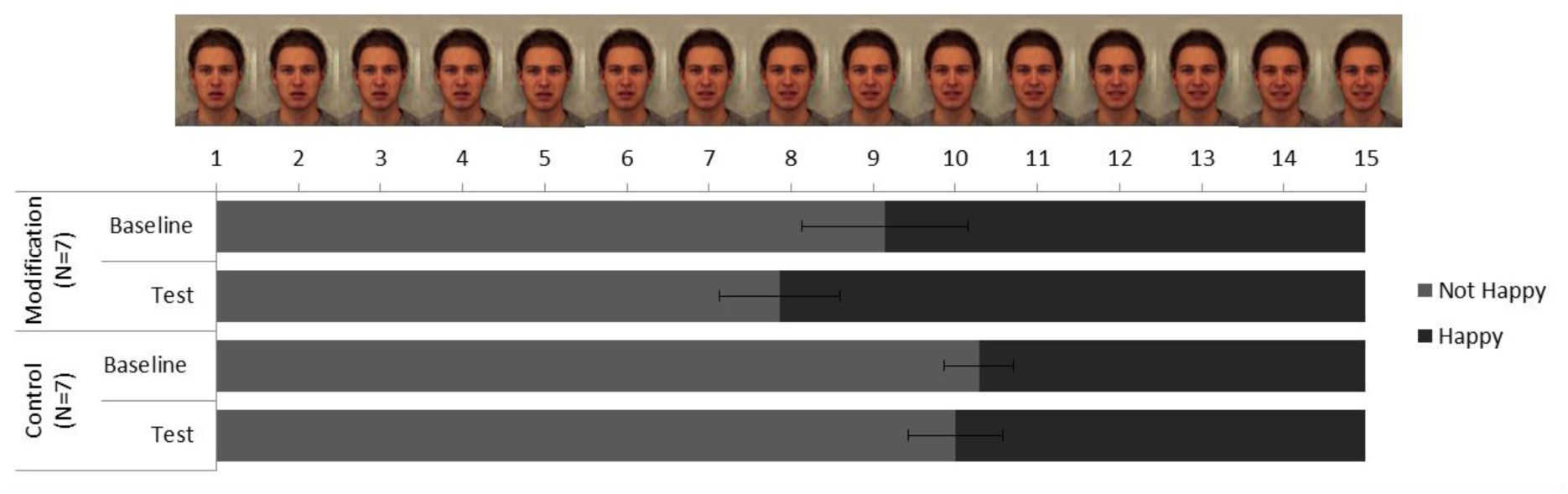
Mean thresholds recorded in baseline and test phases (proportion of “happy” responses) in the modification and control conditions in Study 1. Error bars represent standard errors.

**Figure 2.**
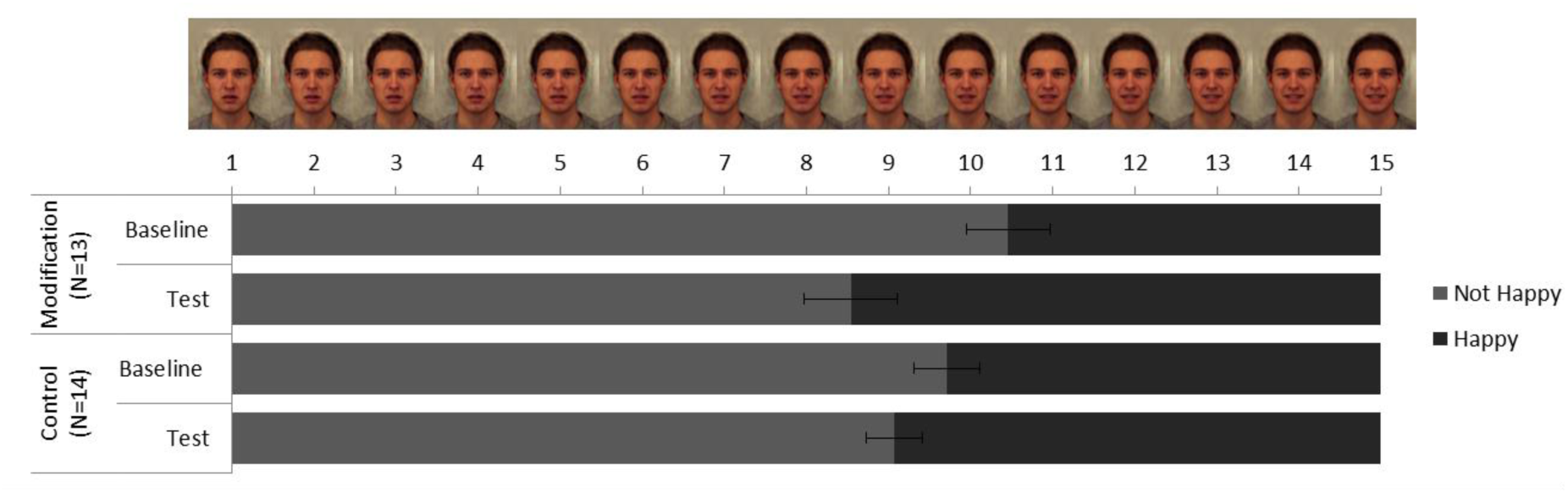
Mean thresholds recorded in baseline and test phases (proportion of “happy” responses) in the modification and control conditions in Study 2. Error bars represent standard errors.

ANOVA revealed evidence of a main effect of time (*F* [1, 12] = 9.55, *p* =.009, η^2^ =.44), reflecting the fact that in both conditions thresholds were lower in the test phase than at baseline. There was no clear evidence for a main effect of condition (*F* [1, 12] = 2.77, *p* =.12, η^2^ =.19), but some evidence of an interaction between time and condition (*F* [1, 12] = 3.87, *p* =.073, η^2^ =.24), Post-hoc t-tests confirmed that this interaction was due to a decrease in threshold from pre to post training in the group which received modification feedback (*t* [6] = 3.06, *p* =.022) but not in the group which received control feedback (*t* [6] = 1.00, *p* =.36).

## Discussion

In this first experiment we found some statistical evidence that participants who received modification feedback showed an increase in the number of expressions from the happy to anti-happy morph sequence that they identified as happy, compared to those who received control feedback. These findings suggest that modification training led to an increase the detection of happiness in these low intensity expressions.

In order to strengthen the evidence that this training is possible in individuals with ASD, the next experiment aimed to replicate the finding of Study 1 in a group of adults with ASD. Self-report measures of the severity of ASD symptoms and the level of depression symptoms were collected. Both ASD and depression (which is common in adults with ASD, Joshi et al., 2010), have been associated with reduced recognition of happy expressions (Bourke et al., 2010; Eack et al., 2015; Humphreys, Minshew, Leonard, & Behrmann, 2007). Greater ASD or depression symptoms may therefore be associated with differences in the malleability of thresholds. In order to account for this we controlled for symptom levels in the analysis. Additionally we looked at correlations between the size of the training effect and level of ASD and depression symptoms to see if these affected the success of training in altering thresholds.

Additionally we tested whether increasing detection of low intensity happiness using morph sequence training improved the accuracy of expression recognition more generally, using a standardised measure of facial expression perception which has been shown to be sensitive to emotion recognition impairments in ASD (Baron-Cohen, Wheelwright, Hill, Raste, & Plumb, 2001). This study was preregistered as a randomised controlled trial (ISRCTN97201297).

## Study 2

### Method

#### Recruitment

Adults (23-63 years) with ASD were recruited through a specialist National Health Service facility (n = 25) and the University of Bristol disability service (n = 3). Confirmation of diagnosis was provided by the NHS service or the University disability service with reference to medical records or service records respectively. This study was approved by the University of Bristol Faculty of Science Research Ethics Committee and the National Health Service Research Ethics Committee.

#### Measures

The feedback training task was as described in Study 1. Reading the Mind in the Eyes task (RME; Baron-Cohen et al., 2001) was used to measure expression perception. RME consists of 36 trials in which participants are shown a photo of the eye region and chose one of four complex mental state (e.g. confused, ashamed, worried) to match the photo. The Ritvo Autism Asperger’ s Diagnostic Scale-Revised (RAADS-R; Ritvo et al., 2011) was used to measure ASD symptoms. RAADS-R is an 80 item questionnaire for measuring ASD traits in adults. Finally, the 21 item Beck Depression Inventory questionnaire (BDI-II; Beck, Steer & Brown, 1996) was used to measure depressive symptoms.

#### Procedure

Testing took place either at the participant’s home, on site at the National Health Service facility or community advice service, or at the University of Bristol. Participants first completed the three phases of the training task as described in Study 1. All participants pressed the response keys themselves. The RME task was completed after the training task. The BDI and the RAADS-R questionnaires were either completed before coming to the session or completed after the training task.

#### Analysis

Thresholds were analysed with a mixed model ANOVA as in Study 1. This was done both with and without BDI-II and RAADS-R scores included as covariates. Correlations between change in threshold and BDI-II and RAADS-R scores were analysed to determine whether symptom severity or depression symptoms were associated with change in threshold.

## Results

### Participants

A total of twenty-eight participants were recruited. One participant did not complete the computer task, leaving a total of 27 participants, 13 who had been randomly assigned to the modification condition (2 females, mean age = 37 years, SD = 15 years), and 14 who had been randomly assigned to the control condition (1 female, mean age = 40 years, SD = 12 years). Twenty three participants completed the RAADS-R and all scored well above the cut-off for ASD (mean score 141.32, range 90-195, cut-off 65). Five participants did not complete this measure due to time constraints.

### Training effects

The mean thresholds at baseline were 10.46 (*SD* = 1.85) frames in the modification condition and 9.71 (*SD* = 1.49) frames in the control condition. The mean threshold at test was 8.54 (*SD* = 2.03) frames in the modification condition and 9.07 (*SD* = 1.27) frames in the control condition (see Figure 2).

ANOVA revealed evidence of a main effect of time (*F* [1, 25] =17.86, *p* <.001, η^2^=.42). There was no evidence for a main effect of condition (*F* [1, 25] = 0.04, *p* =.85, η^2^=.001), but there was evidence of an interaction between time and condition (*F* [1, 25] = 4.45, *p* =.045, η^2^=.15), reflecting the greater decrease in threshold from baseline to test in the modification condition compared to the control condition (see Figure 3). When BDI-II and RAADS-R scores included as covariates, the evidence for the critical interaction between time and condition remained, although the statistical evidence was weaker (*F* [1, 25] = 3.63, *p* =.069).

Post-hoc t-tests confirmed that the interaction between time and condition was due to a decrease in threshold in the modification condition (*t* [12] = 5.84, *p* <.001) but not the control condition (*t* [13] = 1.29, *p =*.22) in the unadjusted analysis, but again strength of evidence was attenuated when BDI-II and RAADS-R were included as covariates (modification *p* =.12, control *p* = 1.0).

The BDI-II was completed by 26 participants (modification: mean = 17.96, *SD* = 8.04; control: mean = 24.32, *SD* = 14.72), while the RAADS-R was completed by 23 participants (modification: mean = 137.50, *SD* = 25.04; control: mean = 145.14, *SD* = 27.94). Not all participants completed these measures due to time constraints in the testing session. There was no meaningful correlation between change in threshold and either BDI-II scores (*r* [24] = -.15, *p* =.47) or RAADS-R scores (*r* [21] = -.05, *p* =.82).

Finally, there was no evidence that the group that received modification training performed differently to the group that received control training on the RME task completed after training (modification: mean = 24.64, *SD* = 4.96; control: mean = 21.92, *SD* = 7.11; *t* [25] = 1.16, *p* =.26).

## Discussion

Study 2 replicated the findings of Study 1 in a population of adults with ASD. As in Study 1, there was some evidence that participants who received modification feedback showed an increase in the number of expressions from the happy to anti-happy morph sequence that they identified as happy, compared to those who received control feedback. The fact that the findings of Study 1 were replicated in a preregistered study adds strength to the conclusion that the intended training effects can be achieved in populations with ASD.

Self-report measures of ASD and depression symptoms were taken to control for any differences in these variables that existed between groups. There was no statistical evidence that ASD symptom severity nor depression symptom severity correlated with the size of training effects, indicating that variation in level of ASD and depression does not have a substantial influence on the effect of training. However, the control group did have higher levels of depression and ASD symptoms than the modification group, and when depression and ASD symptom severity were controlled for in the analysis of threshold change, evidence for the group difference attenuated. This may suggest that some of the group difference in threshold change was the result of higher levels of symptom severity in the control group reducing the potential for threshold change in this group compared to the modification group. Alternatively, the attenuation of the effect with the addition of the covariates may be the result of reduced statistical power to find a group difference in the analysis.

We found no difference in performance on the RME task between groups after training. Assuming that the two groups would have performed similarly on this test at baseline, the lack of group difference in performance after training suggests that the training procedure does not have an immediate impact on the accuracy of global facial expression recognition. However, in the absence of baseline measurement it is not possible to conclude that there was no effect of the training on performance, as one group may have started off performing more poorly and have improved more.

## General Discussion

The two studies reported here pilot test a training methodology which aims to increase the recognition of happiness in low intensity happy expressions. The training paradigm was originally designed to address biases in emotion perception observed in depression or conduct disorder (Penton-Voak et al., 2012; Penton-Voak et al., 2013). For these populations the training aimed to reduce biases towards perceiving sadness or anger, by encouraging perception of happiness in expressions taken from morph sequences morphing between happy and angry, or happy and sad expressions. We modified this paradigm in an attempt to increase detection of happiness in low intensity happy expressions without decreasing detection of any other emotion. This was done using a morph sequence that mixed a happy expression with a norm expression to create a sequence of varying intensity happy expressions. This modified feedback training method was successful in increasing the number of expressions from the morph sequence that individuals with ASD identified as happy. However, there was no evidence that the feedback training increased facial expression recognition accuracy, as measured by the RME task (Baron-Cohen et al., 2001). While it is possible that the training may have had a specific effect on accuracy of detection of happy expressions, which would not have been picked up by the RME test, our more recent published work suggests that this is not the case (Griffiths, Jarrold, Penton-Voak, & Munafò, 2015). In a study with typically developing adults we found that the same training paradigm used in this study did not increase accuracy of emotion recognition, but rather induces a bias in recognition. This training may therefore be useful for addressing negative interpretation biases in emotion recognition, but will not be useful for increasing accurate detection of emotional expressions, and may actually reduce global recognition accuracy.

